# Global expansion and redistribution of Aedes-borne virus transmission risk with climate change

**DOI:** 10.1101/172221

**Authors:** Sadie J. Ryan, Colin J. Carlson, Erin A. Mordecai, Leah R. Johnson

## Abstract

Forecasting the impacts of climate change on Aedes-borne viruses—especially dengue, chikungunya, and Zika—is a key component of public health preparedness. We apply an empirically parameterized model of viral transmission by the vectors *Aedes aegypti* and *Ae. albopictus*, as a function of temperature, to predict cumulative monthly global transmission risk in current climates, and compare them with projected risk in 2050 and 2080 based on general circulation models (GCMs). Our results show that if mosquito range shifts track optimal temperature ranges for transmission (21.3 – 34.0°C for *Ae. aegypti;* 19.9 – 29.4°C for *Ae. albopictus)*, we can expect poleward shifts in Aedes-borne virus distributions. However, the differing thermal niches of the two vectors produce different patterns of shifts under climate change. More severe climate change scenarios produce larger population exposures to transmission by *Ae. aegypti*, but not by *Ae. albopictus* in the most extreme cases. Climate-driven risk of transmission from both mosquitoes will increase substantially, even in the short term, for most of Europe. In contrast, significant reductions in climate suitability are expected for *Ae. albopictus*, most noticeably in southeast Asia and west Africa. Within the next century, nearly a billion people are threatened with new exposure to virus transmission by both *Aedes* spp. in the worst-case scenario. As major net losses in year-round transmission risk are predicted for *Ae. albopictus*, we project a global shift towards more seasonal risk across regions. Many other complicating factors (like mosquito range limits and viral evolution) exist, but overall our results indicate that while climate change will lead to increased net and new exposures to Aedes-borne viruses, the most extreme increases in *Ae. albopictus* transmission are predicted to occur at intermediate climate change scenarios.

**Author Summary:** The established scientific consensus indicates that climate change will severely exacerbate the risk and burden of Aedes-transmitted viruses, including dengue, chikungunya, Zika, and other significant threats to global health security. Here, we show more subtle impacts of climate change on transmission, caused primarily by differences between the more heat-tolerant *Aedes aegypti* and the more heat-limited *Ae. albopictus.* Within the next century, nearly a billion people could face their first exposure to viral transmission from either mosquito in the worst-case scenario, mainly in Europe and high-elevation tropical and subtropical regions. However, while year-round transmission potential from *Ae. aegypti* is likely to expand (particularly in south Asia and sub-Saharan Africa), *Ae. albopictus* transmission potential is likely to decline substantially in the tropics, marking a global shift towards seasonal risk as the tropics eventually become too hot for transmission by *Ae. albopictus.* Complete mitigation of climate change to a pre-industrial baseline may protect almost a billion people from arbovirus range expansions; however, middle-of-the-road mitigation could produce the greatest expansion in the potential for viral transmission by *Ae. albopictus*. In any scenario, mitigating climate change would shift the projected burden of both dengue and chikungunya (and potentially other *Aedes* transmitted viruses) from higher-income regions back onto the tropics, where transmission might otherwise begin to decline due to rising temperatures.

## Introduction

Climate change will have a profound effect on the global distribution and burden of infectious diseases [1–3]. Current knowledge suggests that the range of mosquito-borne diseases could expand dramatically in response to climate change [4,5]. However, the physiological and epidemiological relationships between mosquito vectors and the environment are complex and often nonlinear, and experimental work has shown a corresponding nonlinear relationship between warming temperatures and disease transmission [6–8]. In addition, pathogens can be vectored by related species, which may be sympatric, or several pathogens may be transmitted by the same vector. Accurately forecasting the potential impacts of climate change on *Aedes-borne* viruses—which include widespread threats like dengue and yellow fever, as well as several emerging threats like chikungunya, Zika, West Nile, and Japanese encephalitis—thus becomes a key problem for public health preparedness [4, 9,10]. In this paper, we compare the impact of climate change on transmission by two vectors, *Aedes aegypti* and *Ae. albopictus.*

The intensification and expansion of vector-borne disease is likely to be a significant threat posed by climate change to human health [2,11]. Mosquito vectors are of special concern due to the global morbidity and mortality from diseases like malaria and dengue fever, as well as the prominent public health crises caused by several recently-emergent viral diseases like West Nile, chikungunya, and Zika. The relationship between climate change and mosquito-borne disease is perhaps best studied, in both experimental and modeling work, for malaria and its associated *Anopheles* vectors. While climate change could exacerbate the burden of malaria at local scales, more recent evidence challenges the “warmer-sicker world” expectation [12,13]. The optimal temperature for malaria transmission has recently been demonstrated to be much lower than previously expected [14], likely leading to net decreases and geographic shifts in optimal habitat at continental scales in the coming decades [13].

Relative to malaria, less is known about the net impact of climate change on Aedes-borne diseases and their vectors. *Ae. aegypti* and *Ae. albopictus* have both (re)emerged and spread worldwide throughout tropical, sub-tropical, and temperate zones in urban areas. *Ae. aegypti* is restricted to warm, urban environments where it breeds in human-made containers in and around houses. Further, it bites during the daytime exclusively on humans, and is a highly competent vector for dengue, chikungunya, Zika, yellow fever, and other viruses. In contrast, while *Ae. albopictus* can adopt this urban ecology, it is far more ecologically flexible, and occurs in suburban, rural, residential, and agricultural habitats and breeds in both natural or human-made containers. It bites humans and a wide range of mammals and birds, and is a moderately competent vector for dengue, chikungunya, and Zika.

In a changing climate, differences in the thermal tolerances of the two mosquitoes are likely to have broad repercussions for their role in future outbreaks. *Ae. albopictus* has become established across a broad latitudinal gradient, with temperate populations undergoing diapause to survive cold winters, while populations in warmer locations have lost the ability to diapause [15]. Aligning with the warmer microclimates of urban environments, *Ae. aegypti* is estimated to have a higher thermal optimum for transmission than *Ae. albopictus* (29°C vs. 26°C) [6]. Given the climate limitations on vector distributions, at a minimum *Aedes* mosquitoes are projected to shift geographically and seasonally in the face of climate change, with a mix of expansions in some regions and contractions in others, and no overwhelming net global pattern of gains or losses [3,9]. Ecophysiological differences between *Aedes* vector species are likely to drive differences in thermal niches, and therefore different distributions of transmission risk [6,16], now and in the future. The consequences of those range shifts for disease burden are therefore likely to be important but challenging to summarize across landscapes and pathogens.

Dengue has reemerged since the 1980s and is now one of the most common vector-borne diseases worldwide, following the end of decades of successful control of the *Ae. aegypti* vector [17,18]. Of all Aedes-borne diseases, dengue fever has been most frequently modeled in the context of climate change, and several models of the potential future of dengue have been published over the last two decades, with some limited work building consensus among them [4]. Models relating temperature to dengue vectorial capacity (the number of new infectious mosquito bites generated by one human case), and applying general circulation models (GCMs) to predict the impacts of climate change, date back to the late 1990s [5]. A study from 2002 estimated that the population at risk (PAR) from dengue would rise from 1.5 billion in 1990 to 5-6 billion by 2085 as a result of climate change [19]. A more recent study added GDP per capita as a predictor in dengue distribution models, and found that climate change would increase the global dengue PAR by a much more moderate 0.28 billion by 2050 with GDP held constant compared to today [20]. Accounting for expected changes in global economic development using linked GDP-climate pathways further reduced the projected PAR by 0.12 billion over the same interval [20]. Mechanistic models have shown that increases or decreases in dengue risk can be predicted for several sites on the same continent based on climate model and scenario selection [21]. Most recently, a recent study prepared for the IPCC 1.5° report showed that limiting climate change to 1.5° could prevent 3.3 million dengue cases per year in the Americas compared to no intervention (+3.7°C) [22].

Chikungunya and Zika viruses, which have emerged more recently as public health crises, are less well-studied in the context of climate change. Like dengue, these viruses are transmitted primarily by *Ae. aegypti* in the tropics and sub-tropics and by *Ae. albopictus* in temperate zones and other settings where *Ae. aegypti* are absent [23–27]. While dengue, chikungunya, and Zika viruses all have sylvatic cycles involving forest mosquitoes and nonhuman primates, recent global outbreaks have been dominated by urban transmission by *Ae. aegypti* and *Ae. albopictus.* Because these viruses are transmitted by the same vectors, and vector physiology dominates much of the transmission process, most early models of climate-dependent chikungunya and Zika transmission assumed that these viruses would exhibit similar thermal responses to dengue [6,28]. However, several virus-specific transmission models have been developed recently. A monthly model for chikungunya in Europe, constrained by the presence of *Ae. albopictus*, found that the A1B and B1 scenarios (older climate change scenarios, roughly comparable to intermediate scenarios RCP 6.0 and 4.5 in current climate assessments) both correspond to substantial increases in chikungunya risk surrounding the Mediterranean [29]. An ecological niche modeling study conducted early in the Zika epidemic found that dengue is likely to expand far more significantly due to climate change than Zika [10] (but epidemiological differences among these three viruses remain unresolved [30–32]). While the combined role of climate change and El Niño has been posited as a possible driver of the Zika pandemic’s severity [10], there is little evidence that anomalous climate conditions were a necessary precursor for Zika transmission. The climate suitability of the region was present and adequate for outbreaks of dengue and chikungunya, transmitted by the same mosquitoes, making the introduction of another arbovirus plausible. In contrast to statistical models, global mechanistic forecasts accounting for climate change are scarce for both chikungunya and Zika, given how recently both emerged as public health crises, and how much critical information is still lacking in the basic biology and epidemiology of both pathogens.

In this study, we apply a new mechanistic model of the spatiotemporal distribution of Aedes-borne viral outbreaks to resolve the role climate change could play in the geographic shifts of diseases like dengue, chikungunya, and Zika. Whereas other mechanistic approaches often rely on methods like dynamic energy budgets to build complex biophysical models for *Aedes* mosquitoes [33,34], and subsequently (sometimes) extrapolate potential epidemiological dynamics [5], our approach uses a single basic cutoff for the thermal interval where viral transmission is possible. The simplicity and transparency of the method masks a sophisticated underlying model that links the basic reproduction number, *R*_0_, for Aedes-borne viruses to temperature, via experimentally-determined physiological response curves for traits like biting rate, fecundity, mosquito lifespan, extrinsic incubation rate, and transmission probability [6].

The models examine the relative sensitivity of *R*_0_ to temperature, conditioned on the presence of the mosquito vector and independent of other factors that might influence transmission, including precipitation, vector control, and prior immune history. The temperature-dependent *R*_0_ model is easily projected into geographic space by defining model-based measures of suitability and classifying each location in space as suitable or not. We parameterize the model using a Bayesian approach to account for uncertainty in the experimental data. The threshold condition in our model (*R*_0_(*T*) > 0) defines the temperatures at which transmission is not prevented, rather than the more familiar threshold at which disease invasion is expected (*R*_0_(*T*) > 1, which cannot be predicted in the absence of assumptions about vector and human population sizes and other factors). We then classify each location by suitability in each month based on already published projections under current climate in the Americas [6].

Here, we extend previous work to investigate the impacts of climate change on Aedes-borne virus transmission. Specifically, we expand the framework for both *Ae. aegypti* and *Ae. albopictus* to project cumulative months of suitability in current and future (2050 and 2080) climates, and further examine how global populations at risk might change in different climate change scenarios. We explore variation among both climate model selection (general circulation models; GCMs), and potential emissions pathways described in the IPCC AR5 (representative concentration pathways; RCPs). In doing so, we provide the first mechanistic forecast for the potential future transmission risk of chikungunya and Zika, which have been forecasted primarily via phenomenological methods (like ecological niche modeling [10]). Our study is also the first to address the seasonal aspects of population at risk for Aedes-borne diseases in a changing climate.

## Methods

### The Temperature Dependent R_0_ Model

Our study presents geographic projections of published, experimentally-derived mechanistic models of viral transmission by *Ae. aegypti* and *Ae. albopictus.* The approach is to fit the thermal responses of all the traits that are components of *R*_0_ in a Bayesian framework and then combine them to obtain the posterior distribution of *R*_0_ as a function of these traits (described in detail in Johnson *et al.* [7], and the particular traits and fits for *Ae. aegypti* and *Ae. albopictus* are presented in Mordecai *et al.* [6]). The approach involves deriving an equation for *R*_0_ from a modified version of the Ross-MacDonald model for mosquito-borne transmission:

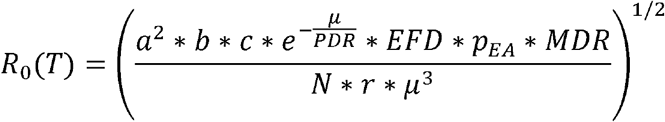

In this equation the parameters are defines as follows: *a* is biting rate (per mosquito); *b*c*is vector competence; *μ* is mosquito mortality rate; *PDR* is parasite (here, viral) development rate; *EFD* is eggs produced per female mosquito per day; *MDR* is the mosquito egg-to-adult development rate; *p_EA_* is probability of mosquito survival from egg to adult; *N* is human population size; and *r* is human recovery rate. All of these parameters, except for *N* and *r*, correspond to vector or pathogen traits and are treated as functions of temperature *T*, with thermal response curves fit to each parameter/trait independently. Because *N* and *r* are difficult to obtain at large scales, *R*_0_ values are rescaled to range from zero to one, and areas where a normalized *R*_0_ > 0 are considered thermally suitable for transmission because temperature does not prevent transmission from occurring. The original study fit trait thermal response curves for each temperature-dependent trait and mosquito species using laboratory experimental data, often derived from single laboratory populations [6]. Further individual- and population-level trait variation remains an empirical gap that is not addressed in the *R*_0_ model.

In the original modeling study [6], empirical data were compiled on transmission of dengue virus by both mosquito species, and the models for *Ae. aegypti* were subsequently validated on human case data compiled for three viruses (dengue, chikungunya, and Zika). The model performed well describing the observed thermal range of transmission for all three viruses, and indicated it was likely adequate for chikungunya and Zika in the absence of more tailored information. Specifically, a statistical model based on the *Ae. aegypti R*_0_(*T*) model predicted the probability of autochthonous transmission in country-scale weekly disease data across the Americas with 86-91% out-of-sample accuracy for dengue and with 66-69% accuracy for chikungunya and Zika, and predicted the magnitude of incidence given local transmission for all three viruses with 85-86% accuracy [6]. A subsequent study using the same *R*_0_(*T*) approach and updated Zika-specific thermal response data has found that the thermal optimum and upper thermal limit for Zika transmission were the same as those of dengue but that the minimum temperature for transmission was higher than that of dengue [35], indicating that the model’s accuracy may vary slightly across viruses transmitted by the focal mosquitoes. Our results are most applicable to dengue, and may offer an indication of possible future scenarios for other viruses, which can be refined as more virus-specific data are collected. For many emerging viruses of concern transmitted by *Aedes* mosquitoes (like Mayaro or St. Louis encephalitis viruses), the data necessary to parameterize temperature-dependent transmission models may not be available for several years.

Once we obtained our posterior samples for scaled *R*_0_ as a function of temperature we evaluated the probability that *R*_0_ > 0 (Prob(*R*_0_ > 0)) at each temperature, giving a distinct curve for each mosquito species. We then defined a cutoff of Prob(*R*_0_ > 0) = *α* to determine our posterior estimate of the probability that temperature is suitable for transmission; here, we use *α* = 0.975. This very high probability allows us to isolate a temperature window for which transmission is almost certainly not excluded; this is a conservative approach designed to minimize Type I error (inclusion of areas not suitable, and prevent overestimation of potential risk). For *Ae. aegypti*, these bounds are 21.3 – 34.0 °C, and for *Ae. albopictus*, 19.9 – 29.4 °C. The prior study added an 8° daily temperature range and incorporated variation in transmission traits within a single day to derive daily average *R*_0_ estimates, for the model validation component [6]; in this study we used the main constant-temperature models derived in [6], as there is both limited empirical laboratory data on the vector-pathogen response to fine-scale variation, and projecting daily thermal range responses so far in the future would add much more uncertainty into our climate forecasts. Our maps of current risk therefore differ slightly from those presented in Figure 4 in [6], but are otherwise produced using the same methodology.

### Current & Future Climates

Current mean monthly temperature data was derived from the WorldClim dataset (www.worldclim.org) [36]. For future climates, we selected four general circulation models (GCMs) that are most commonly used by studies forecasting species distributional shifts, at a set of four representative concentration pathways (RCPs) that account for different global responses to mitigate climate change. These are the Beijing Climate Center Climate System Model (BCC-CSM1.1); the Hadley GCM (HadGEM2-AO and HadGEM2-ES); and the National Center for Atmospheric Research’s Community Climate System Model (CCSM4). Each of these can respectively be forecasted for RCP 2.6, RCP 4.5, RCP 6.0 and RCP 8.5. The scenarios are created to represent standardized cases of how future climate will respond to emissions outputs, ranging from the best-case scenario for mitigation and adaptation (2.6) to the worst-case, business-as-usual fossil fuel emissions scenario (8.5). The scenarios are denoted by numbers (e.g. 2.6, 8.5) corresponding to increased radiation in W/m^2^ by the year 2100, therefore expressing scenarios of increasing severity in the longest term. However, scenarios are nonlinear over time; for example, in 2050, RCP 4.5 is a more severe change than 6.0, as emissions peak mid-century in 4.5 followed by drastic action, whereas emissions rise more slowly to a higher endpoint in 6.0.

Climate model output data for future scenarios were acquired from the research program on Climate Change, Agriculture, and Food Security (CCAFS) web portal (http://ccafs-climate.org/data_spatial_downscaling/), part of the Consultative Group for International Agricultural Research (CGIAR). We used the model outputs created using the delta downscaling method, from the IPCC AR5. For visualizations presented in the main paper (Figure 2a, b), we used the HadGEM2-ES model, the most commonly used GCM. The mechanistic transmission model was projected onto the climate data using the ‘raster’ package in R 3.1.1 (‘raster’[37]). Subsequent visualizations were generated in ArcMap.

### Population at Risk

To quantify a measure of population at risk, comparable between current and future climate scenarios, we used population count data from the Gridded Population of the World, version 4 (GPW4) [26], predicted for the year 2015. We selected this particular population product as it is minimally modeled *a priori*, ensuring that the distribution of population on the earth’s surface has not been predicted by modeled covariates that would also influence our mechanistic vector-borne disease model predictions. These data are derived from most recent census data, globally, at the smallest administrative unit available, then interpolated to produce continuous surface models for the globe for 5-year intervals from 2000-2020. These are then rendered as globally gridded data at 30 arc-seconds; we aggregated these in R to match the climate scenario grids at 5 minute resolution (approximately 10 km^2^ at the equator). We used 2015 population count as our proxy for the current population, and explored future risk relative to the current population counts. This prevents arbitrary demographic model-imposed patterns emerging, possibly obscuring climate-generated change. We note that these count data reflect the disparities in urban and rural patterns appropriately for this type of analysis, highlighting population-dense parts of the globe. Increasing urbanization would likely amplify the patterns we see, as populations increase overall; however, the lack of appropriate population projections at this scale for 30-50 years in the future limits the precision of the forecasts we provide. We thus opted for a most conservative approach. We finally subdivide global populations into geographic and socioeconomic regions as used by the Global Burden of Disease studies (S1 Figure) [39]. We used the ‘fasterize’ R package [40] to convert these regions into rasters with percent (out of 100) coverage at polygon edges. To calculate population at risk on a regional basis, those partial-coverage rasters were multiplied by total population grids.

## Results

The current predicted pattern of temperature suitability based on mean monthly temperatures (**Figure 1**) reproduces the known or projected distributions of Aedes-borne viruses like dengue [41], chikungunya [30], and Zika [10, 43,44] well. For both *Ae. aegypti* and *Ae. albopictus*, most of the tropics is currently optimal for viral transmission year-round, with suitability declining along latitudinal gradients. Many temperate regions currently lacking major *Aedes* vectors, or otherwise considered unsuitable by previous disease distribution models [10, 41,44], are mapped as thermally suitable for up to six months of the year by our model. In these regions where vectors are present, limited outbreaks may only occur when cases are imported from travelers (e.g. in northern Australia, where dengue is not presently endemic but outbreaks occur in suitable regions [17]; or in mid-latitude regions of the United States, where it has been suggested that traveler cases could result in limited autochthonous transmission [31,33]). In total, our model predicts that 6.01 billion people currently live in areas suitable for *Ae. aegypti* transmission at least part of the year (i.e., 1 month or more) and 6.33 billion in areas suitable for *Ae. albopictus* transmission, with significant overlap between these two populations.

**Figure 1.**
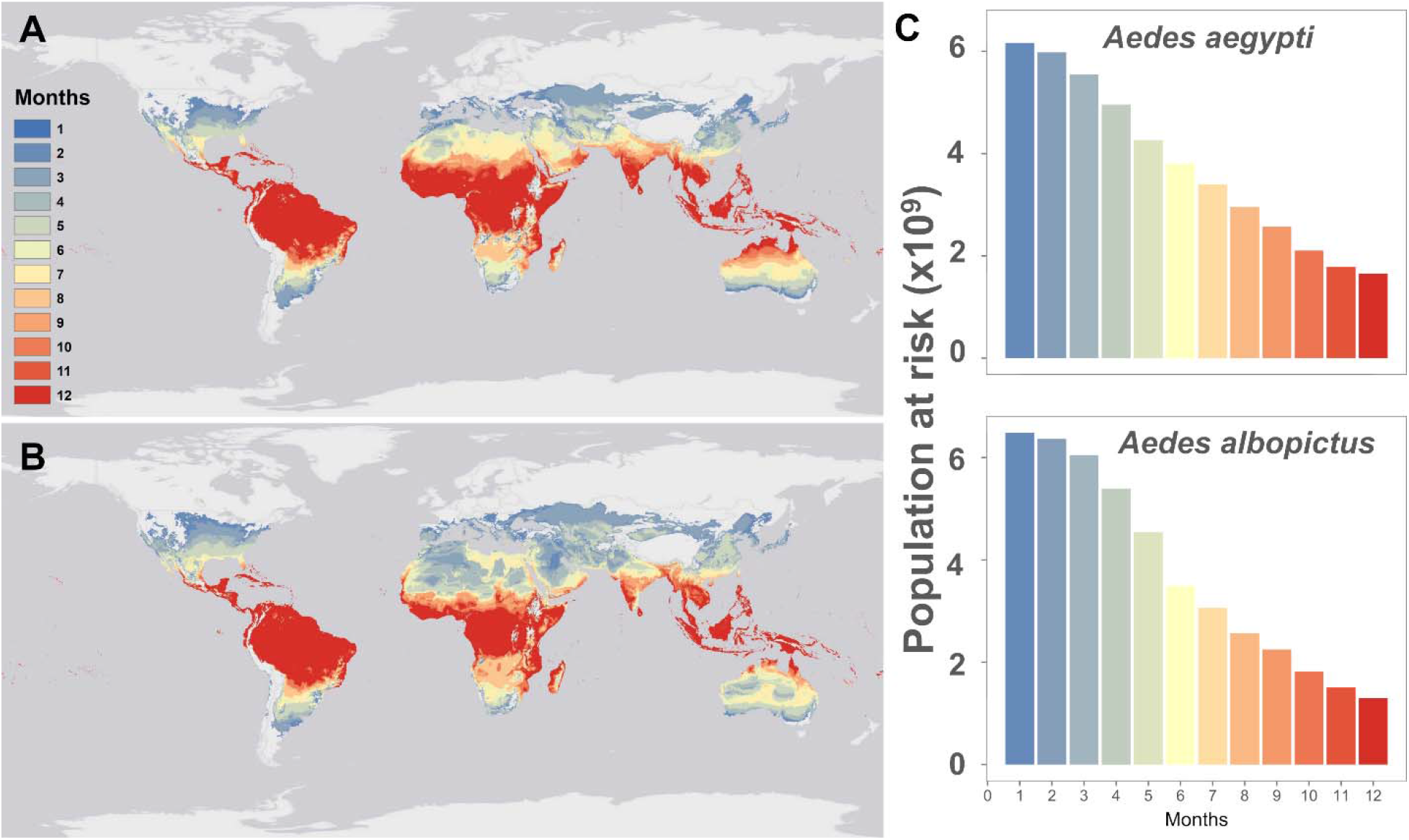
Mapping current temperature suitability for transmission. Maps of current monthly suitability based on mean temperatures using a temperature suitability threshold determined by the posterior probability that scaled *R*_0_ > 0 is 97.5% for (a) *Aedes aegypti* and (b) *Ae. albopictus*, and (c) the number of people at risk (in billions) as a function of their months of exposure for *Ae. aegypti* and *Ae. albopictus.*

By 2050, warming temperatures are expected dramatically expand *Aedes* transmission risk (**Figure 2a, b**). For *Ae. aegypti*, major expansions of one-or two-month transmission risk occur in temperate regions, along with expanding suitability for year-round transmission in the tropics, even into the high-elevation regions that were previously protected by cooler temperatures. *Ae. albopictus* transmission risk similarly expands substantially into temperate regions, especially high latitude parts of Eurasia and North America. However, because warming temperatures are projected to exceed the upper thermal limits to *Ae. albopictus* transmission in many places, major reductions are projected in regions of seasonal risk (e.g., in North Africa) and year-round suitability (e.g., in northern Australia, the Amazon basin, central Africa and southern Asia). Whereas the current gradient of high transmission in the tropics and lower potential in temperate zones is preserved under future climates for *Ae. aegypti*, warming becomes so severe in the tropics that year-round *Ae. albopictus* transmission risk patterns qualitatively change, especially in the more extreme climate pathways. By 2080, year-round temperature suitability for transmission by *Ae. albopictus* is mostly confined to high elevation regions, southern Africa, and the Atlantic coast of Brazil, while the warmer-adapted *Ae. aegypti* begins to lose some core area of year-round temperature suitability for transmission, especially in the Amazon basin.

**Figure 2.**
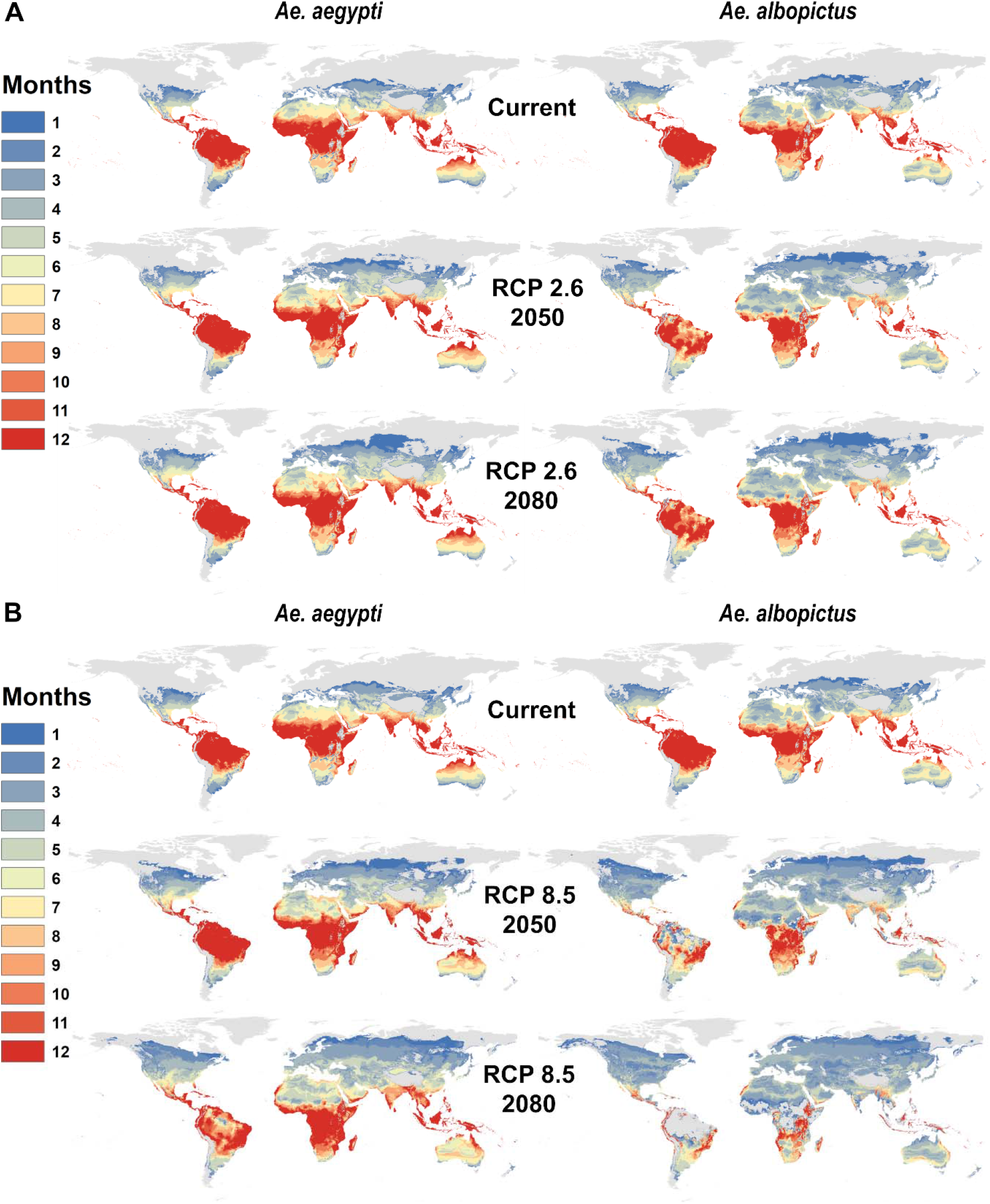
Mapping future temperature suitability for transmission scenarios for *Aedes aegypti and Ae. albopictus.* Maps of monthly suitability based on a temperature threshold corresponding to the posterior probability that scaled *R*_0_ > 0 is greater or equal to 97.5%, for transmission by *Ae. aegypti* and *Ae. albopictus* for predicted mean monthly temperatures under current climate and future scenarios for 2050 and 2080: a. RCP 2.6 and b. RCP 8.5 in HadGEM2-ES.

Concurrently with geographic expansions, our models predict a global net increase in population at risk from Aedes-borne virus exposure, closely tracking the global rise in mean temperatures (**Figure 3**). For both mosquito species, the number of people at risk of ***any months*** of transmission suitability will experience a major net increase by 2050, on the order of half a billion people; however, increases are greater for *Ae. aegypti* than for *Ae. albopictus.* In 2050, more severe climate change scenarios consistently produce greater numbers of people at risk. By 2080, the impact of rising temperature on transmission by each mosquito species diverges: while more severe scenarios continue to drive up the number of people exposed to one or more months of suitable climate for *Ae. aegypti* transmission – to nearly a billion more people exposed than at present – the greatest numbers for *Ae. albopictus* transmission suitability are instead found in intermediate climate change scenarios (RCP 4.5 and 6.0).

**Figure 3.**
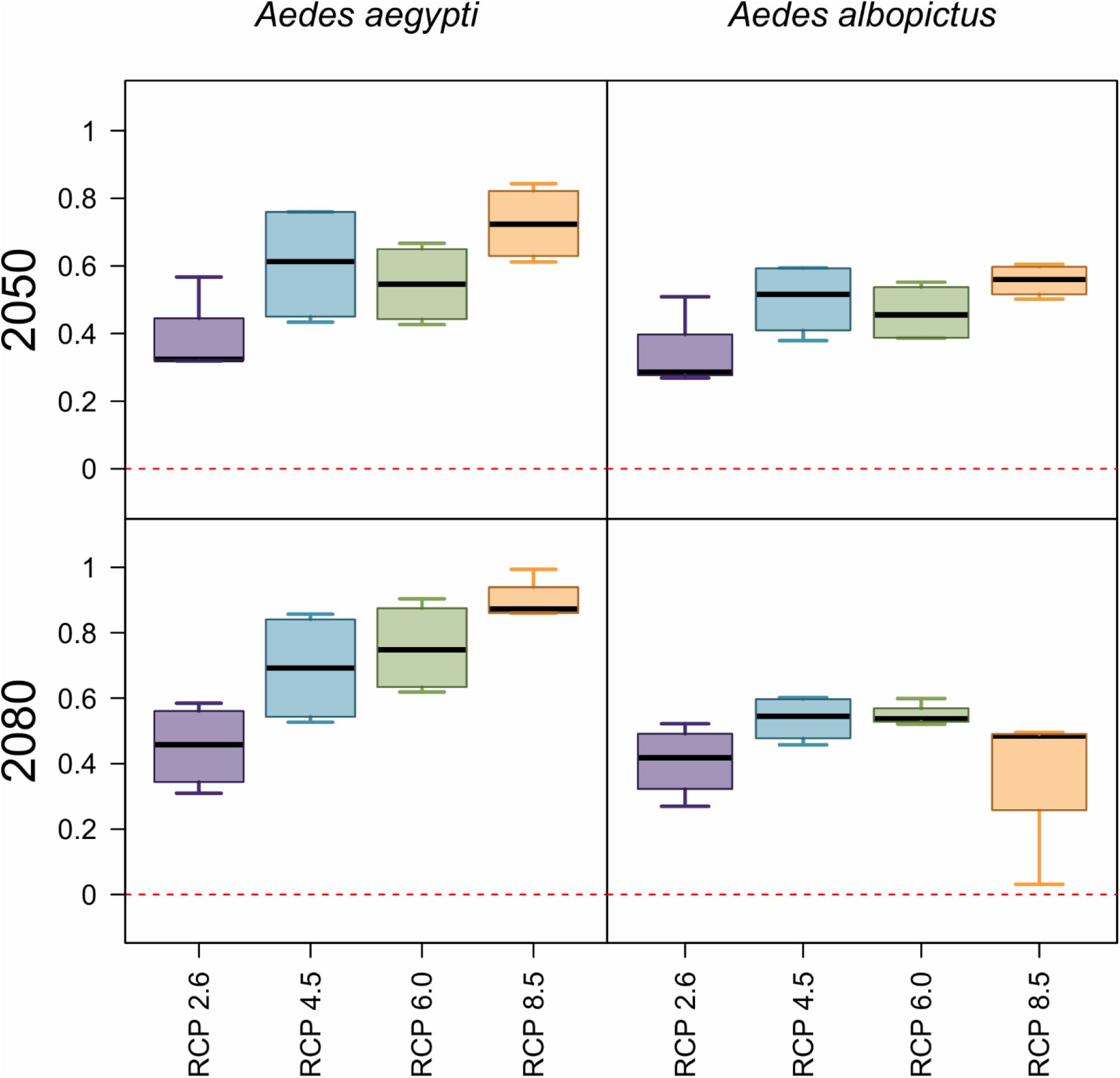
Projected net changes in population at risk. Projections are given as the net difference in billions at risk, for *Ae. aegypti* and *Ae. albopictus* transmission, between current maps and 2050 (top row) or 2080 (bottom row). Results are further broken down by representative climate pathways (RCPs), each averaged across 4 general circulation models.

For year-round exposure, net changes also increasingly differ over time between the two mosquito species. In 2050, warming temperatures lead to a net increase of roughly 100-200 million people in areas of year-round transmission potential for *Ae. aegypti;* in contrast, even in the least severe climate change scenarios, there are drastic net losses of year-round transmission potential for *Ae. albopictus*, and these reductions are larger for more severe scenarios. These patterns continue into 2080 for *Ae. albopictus*, with approximately 700 million fewer people than at present at risk for transmission in the most extreme warming scenarios: in RCP 8.5 by 2080, some parts of the tropics become so warm that even the warmer-adapted *Ae. aegypti* will no longer be able to transmit viruses.

Examining the results by region (**Tables 1 –2**), we find that the regional velocity of climate change is likely to determine the future landscape of *Aedes* transmission risk. For *Ae. aegypti*, increases in the population at risk are expected across the globe and for all climate scenarios and years except in the Caribbean. The most notable net increases in all transmission risk are in Europe, east Asia, high-elevation parts of central America and east Africa, and the United States and Canada. But increases are expected across the board except in the Caribbean, where minor net losses are expected across scenarios and years. In contrast, for *Ae. albopictus*, more region-specific changes are anticipated. Major increases in Europe are expected for all climate scenarios and years, with smaller increases in Central America, east Africa, east Asia, and the U.S. and Canada. However, major net decreases in *Ae. albopictus* transmission potential are expected in several regions, including tropical Latin America, western Africa, south Asia and most of southeast Asia, with a net reduction of nearly 125 million people at risk by 2080 in RCP 8.5. Because the upper thermal limit for *Ae. albopictus* transmission is relatively low (29.4°C), the largest declines in transmission potential in western Africa and southeast Asia are expected with the largest extent of warming, while less severe warming could produce broader increases and more moderate declines in transmission potential. The difference between RCP 6.0 and 8.5 is on the order of 50 million fewer people at risk in west Africa and 100 million fewer in Southeast Asia in the warmer scenario, highlighting just how significant the degree of mitigation will be for regional health pathways.

**Table 1.** Changing population at risk due to temperature suitability for *Aedes aegypti* virus transmission. All values are given in millions; future projections are averaged across GCMs, broken down by year (2050, 2080) and RCP (2.6, 4.5, 6.0, 8.5), and are given as net change from current population at risk. 0+/0- denote the sign of smaller non-zero values that rounded to 0.0, whereas “0” denotes true zeros.

| Region | Current | 2050 |  |  |  | 2080 |  |  |  |
| --- | --- | --- | --- | --- | --- | --- | --- | --- | --- |
|  |  | 2.6 | 4.5 | 6.0 | 8.5 | 2.6 | 4.5 | 6.0 | 8.5 |
| Asia (Central) | 69.9 | 8.4 | 10.5 | 9.9 | 12.2 | 8.1 | 11.8 | 12.5 | 15.6 |
| Asia (East) | 1,321.9 | 42.5 | 49.2 | 46.4 | 58.9 | 38.8 | 56.7 | 61.9 | 72.7 |
| Asia (High Income Pacific) | 164.0 | -0.5 | 0+ | -0.5 | 0.7 | -0.6 | 0.6 | 1.0 | 1.7 |
| Asia (South) | 1,666.4 | -0.1 | 1.6 | 0.7 | 3.7 | -0.5 | 3.4 | 4.3 | 8.2 |
| Asia (Southeast) | 593.9 | -2.1 | 0+ | -0.6 | 2.3 | -2.4 | 1.6 | 2.6 | 5.5 |
| Australasia | 12.9 | 3.6 | 5.7 | 5.3 | 6.7 | 4.3 | 6.2 | 6.9 | 8.0 |
| Caribbean | 40.4 | -1.8 | -1.7 | -1.7 | -1.6 | -1.8 | -1.6 | -1.6 | -1.5 |
| Europe (Central) | 22.7 | 44.2 | 71.8 | 69.0 | 83.3 | 59.0 | 79.3 | 85.5 | 90.6 |
| Europe (Eastern) | 41.3 | 57.9 | 110.4 | 93.5 | 133.9 | 80.0 | 124.7 | 130.7 | 156.2 |
| Europe (Western) | 114.6 | 47.2 | 132 | 112.0 | 166.8 | 90.3 | 156.4 | 180.8 | 220.9 |
| Latin America (Andean) | 31.3 | 2.8 | 3.4 | 3.3 | 4.0 | 2.6 | 3.9 | 4.1 | 5.5 |
| Latin America (Central) | 160.3 | 20.4 | 24.6 | 23.4 | 36 | 18.4 | 34.6 | 39.0 | 61.1 |
| Latin America (Southern) | 42.8 | 8.1 | 8.9 | 8.8 | 9.9 | 7.6 | 9.6 | 10.2 | 12.8 |
| Latin America (Tropical) | 181.8 | 19.2 | 19.5 | 19.5 | 19.6 | 18.9 | 19.6 | 19.7 | 19.8 |
| North Africa & Middle East | 439.5 | 19.7 | 24.1 | 23.8 | 27.2 | 19.3 | 25.9 | 27.3 | 30.3 |
| North America (High Income) | 281.9 | 36.2 | 48.3 | 42.6 | 55.0 | 37.8 | 53.6 | 57.1 | 62.8 |
| Oceania | 6.2 | 0.3 | 0.6 | 0.5 | 0.8 | 0.2 | 0.8 | 0.9 | 1.5 |
| Sub-Saharan Africa (Central) | 115.6 | 5.7 | 6.8 | 6.5 | 7.8 | 5.3 | 7.7 | 8.3 | 9.5 |
| Sub-Saharan Africa (East) | 274.8 | 48.8 | 63.7 | 59.1 | 72.2 | 44.7 | 70.8 | 76.6 | 90.9 |
| Sub-Saharan Africa (Southern) | 46.1 | 23.6 | 25.8 | 25.6 | 26.7 | 23.4 | 26.7 | 27.1 | 28.0 |
| Sub-Saharan Africa (West) | 384.0 | -0.9 | -0.7 | -0.8 | -0.7 | -0.9 | -0.6 | -0.6 | -0.4 |

**Table 2.** Changing population at risk due to temperature suitability for *Aedes albopictus* virus transmission. All values are given in millions; future projections are averaged across GCMs, broken down by year (2050, 2080) and RCP (2.6, 4.5, 6.0, 8.5), and are given as net change from current population at risk. 0+/0- denote the sign of smaller non-zero values that rounded to 0.0, whereas “0” denotes true zeros.

| Region | Current | 2050 |  |  |  | 2080 |  |  |  |
| --- | --- | --- | --- | --- | --- | --- | --- | --- | --- |
|  |  | 2.6 | 4.5 | 6.0 | 8.5 | 2.6 | 4.5 | 6.0 | 8.5 |
| Asia (Central) | 75.7 | 5.0 | 6.9 | 6.4 | 8.8 | 4.7 | 8.1 | 9.1 | 11.2 |
| Asia (East) | 1,367.0 | 16.1 | 20.8 | 18.9 | 25.2 | 15.0 | 24.0 | 26.5 | 32.4 |
| Asia (High Income Pacific) | 167.7 | -2.6 | -2.3 | -2.6 | -2.0 | -2.7 | -2.1 | -1.9 | -2.8 |
| Asia (South) | 1,673.8 | -3.2 | -1.7 | -2.3 | 0+ | -3.5 | -0.5 | -0.3 | -19.1 |
| Asia (Southeast) | 602.5 | -5.3 | -3.8 | -4.0 | -6.7 | -5.4 | -8.5 | -20.1 | -124.8 |
| Australasia | 16.6 | 3.2 | 3.9 | 3.8 | 4.5 | 3.3 | 4.2 | 4.7 | 5.3 |
| Caribbean | 40.8 | -1.8 | -1.8 | -1.8 | -1.8 | -1.9 | -1.8 | -1.8 | -2.3 |
| Europe (Central) | 44.8 | 51.3 | 65.0 | 65.1 | 68.3 | 60.6 | 67.8 | 68.9 | 70.7 |
| Europe (Eastern) | 70.4 | 84.0 | 116.6 | 104.2 | 123.1 | 101.4 | 122.0 | 123.3 | 129.9 |
| Europe (Western) | 135.3 | 98.5 | 179.8 | 161.4 | 208.9 | 149.2 | 199.4 | 215.3 | 243.0 |
| Latin America (Andean) | 33.9 | 1.6 | 2.2 | 2.0 | 2.6 | 1.6 | 2.6 | 2.7 | 2.5 |
| Latin America (Central) | 179.1 | 21.9 | 27.0 | 27.9 | 31.1 | 17.6 | 29.7 | 30.3 | 23.6 |
| Latin America (Southern) | 50.4 | 3.2 | 3.6 | 3.6 | 4.8 | 2.8 | 4.2 | 4.9 | 7.6 |
| Latin America (Tropical) | 203.0 | -1.5 | -2.0 | -1.6 | -6.0 | -1.5 | -5.6 | -8.0 | -26.3 |
| North Africa & Middle East | 455.0 | 10.6 | 13.0 | 12.9 | 14.2 | 10.4 | 13.5 | 14.1 | 11.8 |
| North America (High Income) | 311.6 | 20.6 | 28.4 | 26.0 | 32.1 | 22.6 | 31.6 | 32.3 | 34.7 |
| Oceania | 6.8 | 0.5 | 0.8 | 0.6 | 1.0 | 0.4 | 0.9 | 1.0 | 0.8 |
| Sub-Saharan Africa (Central) | 120.8 | 2.8 | 3.5 | 3.3 | 4.2 | 2.5 | 4.1 | 4.4 | -3.8 |
| Sub-Saharan Africa (East) | 320.2 | 30.3 | 39.1 | 36.3 | 42.4 | 27.9 | 41.8 | 42.8 | 34.2 |
| Sub-Saharan Africa (Southern) | 70.1 | 3.4 | 3.8 | 3.8 | 3.9 | 3.4 | 4.0 | 4.0 | 4.3 |
| Sub-Saharan Africa (West) | 384.9 | -1.4 | -1.5 | -1.5 | -2.0 | -1.4 | -1.9 | -3.5 | -59.0 |

For year-round transmission, the patterns are similarly variable across climate scenarios and regions (**S1–S2 Tables**). Overall, we predict a global shift towards more seasonal risk for both mosquito species, especially in the warmest scenarios. For *Ae. aegypti*, some of the largest net increases in people at risk are expected in southern Africa, with additional notable increases expected in Latin America. Although the upper thermal limit for transmission by *Ae. aegypti* is very high (34.0°C), warming temperatures are projected to exceed the upper thermal limit for parts of the year in some cases; for example, the moderate RCP 4.5 pathway leads to the largest increases in people at risk of temperature suitability for *Ae. aegypti* transmission in southern Asia. Overall, almost 600 million people currently live in areas where temperatures are expected to become suitable for *Ae. aegypti* transmission year-round, though the net increase in year-round transmission will be much less (**S1 Table**). For *Ae. albopictus*, major net reductions are expected in south and southeast Asia, totaling more than 400 million people no longer at year-round risk with the most extreme warming, and additional reductions are expected in east Africa and Latin America. Only the southern part of sub-Saharan Africa is projected to experience net increases in year-round transmission risk (**S2 Table**).

Gross increases—in contrast to net changes—are expected in several regions, particularly in east Africa, placing roughly 250 million people into areas of year-round transmission despite nearly triple that number in net losses. We consider this idea of “first exposures” separately (gross increases, not accounting for losses, in population at any transmission risk), because this form of exposure may be particularly important epidemiologically, as it represents people with little prior immune history with the focal pathogens. While viral immunology is complex [46,47], previously unexposed people may have high susceptibility to chikungunya, Zika, and primary dengue infection, although secondary dengue infections are often the most severe [46]. We rank regions by the number of first exposures (**Table 3**), and we find that consistently the largest number of first exposures driven by newly suitable climate are expected in Europe and east Africa for both mosquito species. As the 2005 epidemic of chikungunya in India and the 2015 pandemic of Zika virus in the Americas highlight, arboviral introductions into naïve populations can produce atypically large outbreaks on the order of millions of infections. This supports concern that both Europe and East Africa may, as a consequence of climate change, be increasingly at risk for explosive outbreaks of vector-borne disease where populations of *Ae. aegypti* and *Ae. albopictus* have established [48,49].

**Table 3.** Top 10 regional increases in overall populations experiencing temperature suitability for transmission (for one or more months). Regions are ranked based on millions of people exposed for the first time to any transmission risk; parentheticals give the net change (first exposures minus populations escaping transmission risk). All values are given for the worst-case scenario (RCP 8.5) in the longest term (2080).

| *Aedes aegypti* |  | *Aedes albopictus* |  |
| --- | --- | --- | --- |
| 1. Europe (Western) | 224 (220.9) | 1. Europe (Western) | 246.2 (243) |
| 2. Europe (Eastern) | 156.4 (156.2) | 2. Europe (Eastern) | 130.1 (129.9) |
| 3. Sub-Saharan Africa (East) | 92.8 (90.9) | 3. Europe (Central) | 71 (70.7) |
| 4. Europe (Central) | 90.9 (90.6) | 4. Sub-Saharan Africa (East) | 58.1 (34.2) |
| 5. Asia (East) | 81.7 (72.7) | 5. Latin America (Central) | 51.9 (23.6) |
| 6. North America (High Income) | 65.7 (62.8) | 6. Asia (East) | 41.4 (32.4) |
| 7. Latin America (Central) | 62 (61.1) | 7. North America (High Income) | 37.7 (34.7) |
| 8. North Africa & Middle East | 34.3 (30.3) | 8. North Africa & Middle East | 19.4 (11.8) |
| 9. Sub-Saharan Africa (Southern) | 28 (28) | 9. Asia (South) | 12.1 (-19.1) |
| 10. Latin America (Tropical) | 21.7 (19.8) | 10. Asia (Central) | 11.2 (11.2) |
| **Total** (across all 21 regions) | **951.3 (899.7)** | **Total** (across all 21 regions) | **721.1 (373.9)** |

## Discussion

The dynamics of mosquito-borne illnesses are climate-driven, and current work suggests that climate change will dramatically increase the potential for expansion and intensification of Aedes-borne virus transmission within the next century. Modeling studies have anticipated climate-driven emergence of dengue and chikungunya at higher latitudes [50,51] and higher elevations [52,53], and predicted the potential ongoing global expansion of Zika [10,44]. The majority of research at global scales [10, 21,54] and in North America and Europe [55] has suggested that climate change is likely to increase the global burden of dengue and chikungunya, and therefore, that mitigation is likely to benefit global health [22,56]. Aedes-borne virus expansion into regions that lack previous exposure is particularly concerning, given the potential for explosive outbreaks when arboviruses are first introduced into naïve populations, like chikungunya and Zika in the Americas [57]. The emergence of a Zika pandemic in the Old World [58], the establishment of chikungunya in Europe beyond small outbreaks [29], or introduction of dengue anywhere a particular serotype has not recently been found, is a critical concern for global health preparedness. However, because effects of climate on vector-borne disease transmission are nonlinear [6,14,35,59–62], climate change may increase transmission potential in some settings and decrease it in others, yet the potential for climate-driven shifts and decreases in disease burden is less well-understood (but see Ryan et al. [13]). Using geospatial projections of a mechanistic temperature-dependent transmission model, we investigated geographic, seasonal, and population changes in risk of virus transmission by two globally important vectors, *Ae. aegypti* and *Ae. albopictus* with climate warming by 2080.

Overall, our findings support the expectation that climate change will expand and increase Aedes-borne viral transmission risk. However, we also find more nuanced patterns that differ among the mosquito species, climate pathways, localities, and time horizons. The largest increases in population at risk are consistently projected in Europe, with additional increases in high altitude regions in the tropics (eastern Africa and the northern Andes) and in the United States and Canada. These increases are expected not only for occasional exposure, but also for longer seasons of transmission, especially by *Ae. aegypti.* However, in the tropics, for both mosquito species, and in other regions for *Ae. albopictus*, more extreme climate pathways are expected to increase temperatures beyond the suitable range for transmission in many parts of the world. In addition to increases in total exposure from both mosquitoes in our study, we predict a global shift towards seasonal regimes of exposure from *Ae. albopictus.* This apparent paradox is due to the shifting geography of two slightly different sets of temperature bounds, as they also move across varying densities of human populations.

As warming temperatures could exceed the upper thermal limits for transmission, estimated at 29.4°C for *Ae. albopictus* and 34.0°C for *Ae. aegypti*, we predict, counterintuitively, that partial climate change mitigation could cause greater increases in exposure risk, particularly for transmission by *Ae. albopictus*, than no mitigation. However, partial mitigation is predicted to decrease the people, geographic areas, and length of seasons of risk for transmission by *Ae. aegypti*, the primary vector of arboviruses like dengue, chikungunya, and Zika, in most regions. Total mitigation (down to pre-industrial baselines) would presumably prevent this redistribution of global risk more effectively. Given the current insufficient response to curb carbon emissions and keep temperatures below the 2°C warming target [63], models such as those presented here can be useful as a means to anticipate possible future changes in climate and temperature-driven transmission risk, depending on the degree of mitigation achieved.

These future predictions of global climate suitability for transmission risk are inherently stochastic, and the degree to which our models will correspond to reality depends not only on uncertainty about climate change, but also on uncertainty about the other environmental, biological, and social factors that drive disease [64]. For example, reductions in transmission may be less prevalent than we expect here, if viruses and vectors evolve higher upper thermal limits for transmission. Because mosquito survival is most limiting to transmission at high temperatures for both mosquito species [6], increasing the thermal limits for transmission would require mosquitoes to adapt to higher survival at warm temperatures, but selective pressure on mosquitoes might instead promote faster development and reproductive cycles over short lifespans. The extent to which mosquitoes and viruses can adapt to warming temperatures remains a critical empirical gap, though there is limited evidence of a mosquito genotype by chikungunya virus strain by temperature interaction in the laboratory, suggesting that populations may vary in transmission potential across temperatures [65]. Increases in transmission risk are also complicated by other factors that drive transmission such as the presence or absence of *Aedes* mosquitoes, which are also undergoing range shifts facilitated by both climate change and human movement. Our model describes areas where *Ae. albopictus* and *Ae. aegypti* are currently absent but could be present in the future, and may represent overestimates of risk should *Aedes* ranges fail to expand into these areas. Whether expanding transmission risk leads to future viral establishment and outbreaks depends not only on pathogen introduction, but also on land use patterns and urbanization at regional scales, which mediate vector distributions and vector – human contact rates [66,67].

In addition, the accuracy of the model predictions for different combinations of vector, virus, and region depends on how vector and virus traits vary across populations and regions. The data used to fit the mechanistic models are derived from dengue virus traits in mosquitoes from multiple source populations from independently-published trait thermal response studies [6]. In addition, many of the traits are measured in laboratory strains and rearing conditions, which may distort life-history relative to wild vector populations, although this will be more problematic for pathogens with longer development times, such as malaria [68]. For *Ae. aegypti*, the most commonly implicated vector of dengue, our results suggest a strong link between warming temperatures and increased transmission [6,41]. The temperature-dependent transmission models were also originally validated for chikungunya and Zika viruses in the Americas and performed well, indicating coarse-scale predictability of climate-dependent patterns of transmission [35]. For chikungunya, the reductions in *Ae. albopictus* transmission potential in south and southeast Asia are particularly notable because the vector is common in that region, where it transmits the introduced Indian Ocean lineage (IOL) of chikungunya (characterized by the E1-226V mutation, which increases transmission efficiency by *Ae. albopictus* specifically [69,70]). In south and southeast Asia, these results might suggest a decreased risk of chikungunya transmission in the most extreme climate scenarios, while arbovirus transmission by *Ae. aegypti* could continue to increase. Further, multiple chikungunya introductions to Europe have been transmitted by *Ae. albopictus* and/or have carried the E1-226V mutation, suggesting that *Ae. albopictus* expansion in Europe might correspond to increased chikungunya risk [69, 71,72]. In contrast, *Ae. aegypti* may be more relevant as a chikungunya vector in the Americas, where it was implicated in the explosive 2015 outbreak [69].

In practice, these models are a first step towards an adequate understanding of potential future changes in the distribution of disease burden, and the potential of these models to make accurate predictions depends on a number of confounding factors [73,74]. In particular, the link from transmission risk to clinical outcomes is confounded by other health impacts of global change, including changing patterns of precipitation, socioeconomic development, nutrition, healthcare access, land use, urbanization, vector and virus adaptation to temperature and other pressures, and vector management, all of which covary strongly. Moreover, human behavioral and societal adaptation to climate change may have just as much of an impact as mitigation in determining how disease risk patterns shift; for example, increased drought stress may encourage water storage practices that increase proximity to *Aedes* breeding habitat [75]. Together concurrent global changes will determine the burden of *Aedes*-borne outbreaks, modulating the predictions we present here.

Many models exist to address this pressing topic, each with different approaches to control for data limitations, confounding processes, climate and disease model uncertainty, different concepts of population at risk, and different preferences towards experimental, mechanistic, or phenomenological approaches e.g. [4,8,10,16,24,33,34,41,44,45,53,61,67,67,76–78]. While climate change poses perhaps the most serious threat to global health security, the relationship between climate change and burdens of Aedes-borne diseases is unlikely to be straightforward, and no single model will accurately predict the complex process of a global regime shift in Aedes-borne viral transmission. Our models only set an outer spatiotemporal bound on where transmission is thermally plausible, given climate projections. Climate change is likely to change the relationship between transmission risk and disease burden at fine scales within those zones of transmission nonlinearly, such that areas with shorter seasons of transmission could still experience increased overall disease burdens, or vice versa. Combining broad spatial models with finer-scale models of attack rates or outbreak size is a critical step towards bridging scales [58,79], but more broadly, building consensus and revealing similarities and differences between all available models via transparency, is of paramount importance [80]. This task is not limited to research on dengue and chikungunya; with several emerging flaviviruses on the horizon [81,82], and countless other emerging arboviruses likely to test the limits of public health infrastructure in coming years [83], approaches that bridge the gap between experimental biology and global forecasting can be one of the foundational methods of anticipating and preparing for the next emerging global health threat.

## Acknowledgements

Van Savage, Naveed Heydari, Jason Rohr, Matthew Thomas, and Marta Shocket provided helpful discussions on modeling approaches.

## Author Information

Correspondence and requests for materials should be addressed to S.J.R.

## Supplementary Figures & Tables

**S1 Figure.**
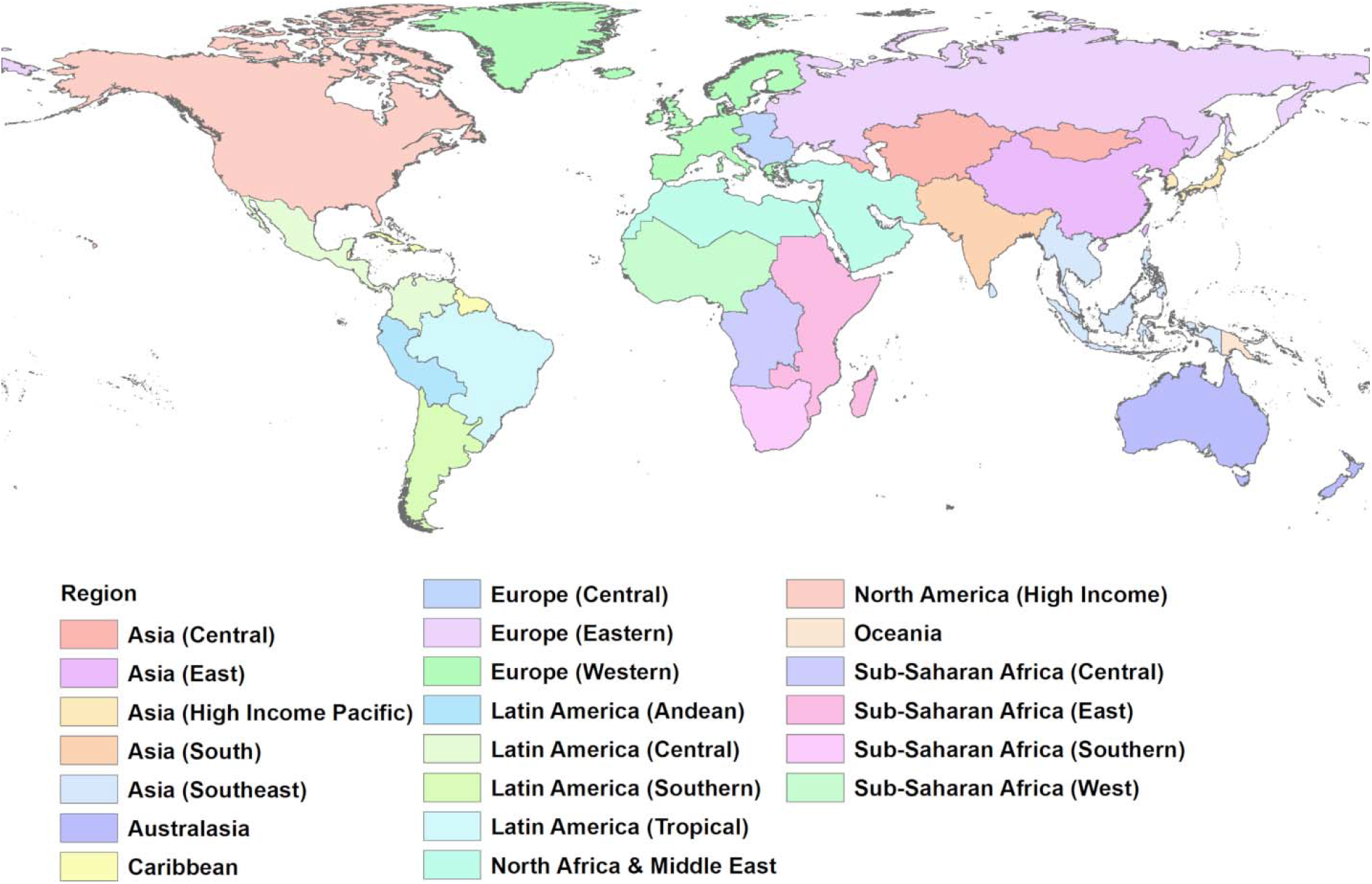
Global health regions. We adopt the same system as the Global Burden of Disease Study in our regional breakdown.

**S1 Table.** Changing year-round (12 month) population at risk due to temperature suitability for *Aedes aegypti* virus transmission. All values are given in millions; future projections are averaged across GCMs, broken down by year (2050, 2080) and RCP (2.6, 4.5, 6.0, 8.5), and are given as net change from current population at risk. 0+/0-denote the sign of smaller non-zero values that rounded to 0.0, whereas “0” denotes true zeros. (Losses do not indicate loss of any transmission, only to reduction 11 or fewer months.).

| Region | Current | 2050 |  |  |  | 2080 |  |  |  |
| --- | --- | --- | --- | --- | --- | --- | --- | --- | --- |
|  |  | 2.6 | 4.5 | 6.0 | 8.5 | 2.6 | 4.5 | 6.0 | 8.5 |
| Asia (Central) | 0 | 0 | 0 | 0 | 0 | 0 | 0 | 0 | 0 |
| Asia (East) | 0+ | 1.4 | 1.8 | 1.1 | 3.7 | 1.6 | 4.5 | 4 | 8.3 |
| Asia (High Income Pacific) | 3.6 | -0.2 | -0.2 | -0.2 | -0.2 | -0.2 | -0.2 | -0.2 | -0.2 |
| Asia (South) | 286.4 | 21.8 | 71.8 | 13.7 | 73.6 | 12.1 | 89.7 | 72.6 | 29.6 |
| Asia (Southeast) | 499.4 | 19.2 | 22.4 | 19.9 | 25.1 | 18.9 | 26.3 | 15.4 | -10.3 |
| Australasia | 0.2 | 0+ | 0+ | 0+ | 0.1 | 0+ | 0.2 | 0.2 | 0.3 |
| Caribbean | 34.8 | 1.8 | 2.2 | 2.1 | 2.8 | 1.7 | 2.6 | 2.9 | 3.3 |
| Europe (Central) | 0 | 0 | 0 | 0 | 0 | 0 | 0 | 0 | 0 |
| Europe (Eastern) | 0 | 0 | 0 | 0 | 0 | 0 | 0 | 0 | 0 |
| Europe (Western) | 0 | 0 | 0 | 0 | 0 | 0 | 0 | 0 | 0+ |
| Latin America (Andean) | 14.0 | 3.9 | 4.8 | 4.6 | 5.7 | 3.5 | 5.4 | 5.8 | 7.5 |
| Latin America (Central) | 88.1 | 13.0 | 18.8 | 17.0 | 25.8 | 12.0 | 24.4 | 27.4 | 34.1 |
| Latin America (Southern) | 0 | 0 | 0 | 0 | 0 | 0 | 0 | 0 | 0.2 |
| Latin America (Tropical) | 67.5 | 27.2 | 34.5 | 30.8 | 41.5 | 27.3 | 39 | 42.9 | 54.9 |
| North Africa & Middle East | 12.5 | -5.2 | -5.5 | -6.0 | -5.6 | -4.7 | -5.4 | -5.4 | -3.9 |
| North America (High Income) | 0.5 | 0.3 | 0.9 | 0.6 | 1.5 | 0.3 | 1.9 | 1.6 | 5.5 |
| Oceania | 0 | 0 | 0 | 0 | 0 | 0 | 0 | 0 | 0 |
| Sub-Saharan Africa (Central) | 5.3 | 0.3 | 0.6 | 0.4 | 0.9 | 0.3 | 0.7 | 0.9 | 1.7 |
| Sub-Saharan Africa (East) | 79.0 | 19.1 | 23.1 | 22.4 | 26.8 | 16.6 | 25.4 | 28.0 | 36.1 |
| Sub-Saharan Africa (Southern) | 126.9 | 43.8 | 60.7 | 56.7 | 78.3 | 37.9 | 74.3 | 85.5 | 110.3 |
| Sub-Saharan Africa (West) | 0 | 0+ | 0.1 | 0.1 | 0.3 | 0+ | 0.2 | 0.6 | 4.4 |

**S2 Table.** Changing year-round (12 month) population at risk due to temperature suitability for *Aedes albopictus* virus transmission. All values are given in millions; future projections are averaged across GCMs, broken down by year (2050, 2080) and RCP (2.6, 4.5, 6.0, 8.5), and are given as net change from current population at risk. 0+/0- denote the sign of smaller non-zero values that rounded to 0.0, whereas “0” denotes true zeros. (Losses do not indicate loss of any transmission, only to reduction 11 or fewer months).

| Region | Current | 2050 |  |  |  | 2080 |  |  |  |
| --- | --- | --- | --- | --- | --- | --- | --- | --- | --- |
|  |  | 2.6 | 4.5 | 6.0 | 8.5 | 2.6 | 4.5 | 6.0 | 8.5 |
| Asia (Central) | 0 | 0 | 0 | 0 | 0 | 0 | 0 | 0 | 0 |
| Asia (East) | 1.3 | 1.4 | -0.4 | -0.5 | -1 | 1 | -0.9 | -1 | -1.2 |
| Asia (High Income Pacific) | 3.6 | -0.3 | -0.5 | -0.4 | -2.9 | -0.2 | -2.2 | -3.1 | -3.6 |
| Asia (South) | 98.3 | -73 | -80.3 | -78.9 | -87.3 | -67.7 | -86.1 | -88.5 | -92.6 |
| Asia (Southeast) | 435.3 | -133.9 | -213.3 | -190.9 | -277.4 | -131.9 | -254.8 | -282.7 | -343.6 |
| Australasia | 0.2 | 0+ | 0+ | 0+ | 0- | 0+ | 0- | 0- | 0- |
| Caribbean | 39.3 | -5.9 | -11.7 | -9.4 | -17.5 | -4.5 | -16.1 | -18.1 | -28.0 |
| Europe (Central) | 0 | 0 | 0 | 0 | 0 | 0 | 0 | 0 | 0 |
| Europe (Eastern) | 0 | 0 | 0 | 0 | 0 | 0 | 0 | 0 | 0 |
| Europe (Western) | 0 | 0 | 0 | 0 | 0+ | 0 | 0+ | 0+ | 0.1 |
| Latin America (Andean) | 17.9 | 2 | 0.1 | 0.5 | -3 | 1.8 | -2 | -3.1 | -5 |
| Latin America (Central) | 97.2 | -23.2 | -26.8 | -25.3 | -31.0 | -20.8 | -29.3 | -31.4 | -33.6 |
| Latin America (Southern) | 0 | 0 | 0 | 0 | 0+ | 0 | 0 | 0+ | 0+ |
| Latin America (Tropical) | 93.6 | -0.8 | -5.2 | -6.1 | -9.7 | -2.9 | -10.1 | -12.9 | -37.1 |
| North Africa & Middle East | 2.6 | 0+ | 0+ | -0.2 | -0.2 | 0.1 | 0- | -0.2 | -0.1 |
| North America (High Income) | 1 | 2.8 | 1.4 | 1 | 0+ | 1.6 | 0.1 | -0.2 | -0.2 |
| Oceania | 0 | 0 | 0 | 0 | 0 | 0 | 0 | 0 | 0 |
| Sub-Saharan Africa (Central) | 5.9 | 0.4 | 0.4 | 0.5 | 0.1 | 0.4 | 0.1 | 0- | -1.4 |
| Sub-Saharan Africa (East) | 96.6 | 8.0 | 5.1 | 6.8 | -4.5 | 7.5 | -9.1 | -9.9 | -45.8 |
| Sub-Saharan Africa (Southern) | 133.5 | 31.9 | 38.8 | 38.8 | 43.4 | 29.7 | 39.5 | 43.9 | 39.2 |
| Sub-Saharan Africa (West) | 0+ | 0+ | 0.5 | 0.4 | 1.8 | 0+ | 0.9 | 2 | 6.2 |

**S3 Table.** Top 10 regional increases in populations experiencing year-round temperature suitability for transmission (12 months). Regions are ranked based on millions of people exposed for the first time to any transmission risk; parentheticals give the net change (first exposures minus populations escaping transmission risk). All values are given for the worst-case scenario (RCP 8.5) in the longest term (2080).

| *Aedes aegypti* |  | *Aedes albopictus* |  |
| --- | --- | --- | --- |
| 1. Asia (South) | 209.9 (29.6) | 1. Sub-Saharan Africa (East) | 114.3 (39.2) |
| 2. Sub-Saharan Africa (East) | 152.6 (110.3) | 2. Latin America (Tropical) | 39.7 (-37.1) |
| 3. Latin America (Tropical) | 63.2 (54.9) | 3. Latin America (Central) | 38.1 (-33.6) |
| 4. Asia (Southeast) | 44 (-10.3) | 4. Sub-Saharan Africa (Central) | 23.1 (-45.8) |
| 5. Latin America (Central) | 40.7 (34.1) | 5. Asia (Southeast) | 16.3 (-343.6) |
| 6. Sub-Saharan Africa (Central) | 36.6 (36.1) | 6. Latin America (Andean) | 8.6 (-5) |
| 7. Sub-Saharan Africa (West) | 8.7 (-130.2) | 7. Sub-Saharan Africa (Southern) | 6.2 (6.2) |
| 8. Asia (East) | 8.3 (8.3) | 8. Sub-Saharan Africa (West) | 2.5 (-194) |
| 9. Latin America (Andean) | 8 (7.5) | 9. North Africa & Middle East | 2.4 (-0.1) |
| 10. North Africa & Middle East | 7.4 (-3.9) | 10. Oceania | 2 (-1.4) |
| **Total** (across all 21 regions) | **597.2 (151.6)** | **Total** (across all 21 regions) | **256.5 (-740.8)** |

## References

1. Hoberg EP, Brooks DR. Evolution in action: climate change, biodiversity dynamics and emerging infectious disease. Phil Trans R Soc B. 2015;370: 20130553.

2. Lafferty KD. The ecology of climate change and infectious diseases. Ecology. 2009;90: 888–900.

3. Escobar LE, Romero-Alvarez D, Leon R, Lepe-Lopez MA, Craft ME, Borbor-Cordova MJ, et al. Declining Prevalence of Disease Vectors Under Climate Change. Sci Rep. 2016;6.

4. Messina JP, Brady OJ, Pigott DM, Golding N, Kraemer MU, Scott TW, et al. The many projected futures of dengue. Nat Rev Microbiol. 2015;13: 230–239.

5. Patz JA, Martens W, Focks DA, Jetten TH. Dengue fever epidemic potential as projected by general circulation models of global climate change. Environ Health Perspect. 1998;106: 147.

6. Mordecai E, Cohen J, Evans MV, Gudapati P, Johnson LR, Lippi CA, et al. Detecting the impact of temperature on transmission of Zika, dengue, and chikungunya using mechanistic models. PLoS Negl Trop Dis. 2017;11: e0005568.

7. Johnson LR, Ben-Horin T, Lafferty KD, McNally A, Mordecai E, Paaijmans KP, et al. Understanding uncertainty in temperature effects on vector-borne disease: a Bayesian approach. Ecology. 2015;96: 203–213.

8. Lowe R, Gasparrini A, Van Meerbeeck CJ, Lippi CA, Mahon R, Trotman AR, et al. Nonlinear and delayed impacts of climate on dengue risk in Barbados: A modelling study. PLOS Med. 2018;15: e1002613. doi:10.1371/journal.pmed.1002613

9. Campbell LP, Luther C, Moo-Llanes D, Ramsey JM, Danis-Lozano R, Peterson AT. Climate change influences on global distributions of dengue and chikungunya virus vectors. Phil Trans R Soc B. 2015;370: 20140135.

10. Carlson CJ, Dougherty ER, Getz W. An ecological assessment of the pandemic threat of Zika virus. PLoS Negl Trop Dis. 2016;10: e0004968.

11. Githeko AK, Lindsay SW, Confalonieri UE, Patz JA. Climate change and vector-borne diseases: a regional analysis. Bull World Health Organ. 2000;78: 1136–1147.

12. Ibelings B, Gsell A, Mooij W, Van Donk E, Van Den Wyngaert S, Domis D, et al. Chytrid infections and diatom spring blooms: paradoxical effects of climate warming on fungal epidemics in lakes. Freshw Biol. 2011;56: 754–766.

13. Ryan SJ, McNally A, Johnson LR, Mordecai EA, Ben-Horin T, Paaijmans K, et al. Mapping physiological suitability limits for malaria in Africa under climate change. Vector-Borne Zoonotic Dis. 2015;15: 718–725.

14. Mordecai EA, Paaijmans KP, Johnson LR, Balzer C, Ben-Horin T, Moore E, et al. Optimal temperature for malaria transmission is dramatically lower than previously predicted. Ecol Lett. 2013;16: 22–30.

15. Lounibos LP, Escher RL, Lourenço-De-Oliveira R. Asymmetric Evolution of Photoperiodic Diapause in Temperate and Tropical Invasive Populations of Aedes albopictus (Diptera: Culicidae). Ann Entomol Soc Am. 2003;96: 512–518. doi:10.1603/0013-8746(2003)096[0512:AEOPDI]2.0.CO;2

16. Brady OJ, Golding N, Pigott DM, Kraemer MU, Messina JP, Reiner Jr RC, et al. Global temperature constraints on Aedes aegypti and Ae. albopictus persistence and competence for dengue virus transmission. Parasit Vectors. 2014;7: 338.

17. Gubler DJ. The Global Threat of Emergent/Re-emergent Vector-Borne Diseases. In: Atkinson PW, editor. Vector Biology, Ecology and Control. Dordrecht: Springer Netherlands; 2010. pp. 39–62. doi:10.1007/978-90-481-2458-9_4

18. Gubler DJ. The Global Emergence/Resurgence of Arboviral Diseases As Public Health Problems. Arch Med Res. 2002;33: 330–342. doi:10.1016/S0188-4409(02)00378-8

19. Hales S, De Wet N, Maindonald J, Woodward A. Potential effect of population and climate changes on global distribution of dengue fever: an empirical model. The Lancet. 2002;360: 830–834.

20. Åström C, Rocklöv J, Hales S, Béguin A, Louis V, Sauerborn R. Potential distribution of dengue fever under scenarios of climate change and economic development. Ecohealth. 2012;9: 448–454.

21. Williams C, Mincham G, Faddy H, Viennet E, Ritchie S, Harley D. Projections of increased and decreased dengue incidence under climate change. Epidemiol Infect. 2016; 1–10.

22. Colón-González FJ, Harris I, Osborn TJ, São Bernardo CS, Peres CA, Hunter PR, et al. Limiting global-mean temperature increase to 1.5–2° C could reduce the incidence and spatial spread of dengue fever in Latin America. Proc Natl Acad Sci. 2018;115: 6243–6248.

23. Mayer SV, Tesh RB, Vasilakis N. The emergence of arthropod-borne viral diseases: A global prospective on dengue, chikungunya and zika fevers. Acta Trop. 2017;166: 155–163. doi:10.1016/j.actatropica.2016.11.020

24. Grard G, Caron M, Mombo IM, Nkoghe D, Mboui Ondo S, Jiolle D, et al. Zika Virus in Gabon (Central Africa) – 2007: A New Threat from Aedes albopictus? PLoS Negl Trop Dis. 2014;8: e2681. doi:10.1371/journal.pntd.0002681

25. Fauci AS, Morens DM. Zika Virus in the Americas — Yet Another Arbovirus Threat. N Engl J Med. 2016;374: 601–604. doi:10.1056/NEJMp1600297

26. Burt FJ, Rolph MS, Rulli NE, Mahalingam S, Heise MT. Chikungunya: a re-emerging virus. The Lancet. 2012;379: 662–671. doi:10.1016/S0140-6736(11)60281-X

27. Leparc-Goffart I, Nougairede A, Cassadou S, Prat C, de Lamballerie X. Chikungunya in the Americas. The Lancet. 2014;383: 514. doi:10.1016/S0140-6736(14)60185-9

28. Perkins TA, Metcalf CJE, Grenfell BT, Tatem AJ. Estimating Drivers of Autochthonous Transmission of Chikungunya Virus in its Invasion of the Americas. PLoS Curr. 2015;7. doi:10.1371/currents.outbreaks.a4c7b6ac10e0420b1788c9767946d1fc

29. Fischer D, Thomas SM, Suk JE, Sudre B, Hess A, Tjaden NB, et al. Climate change effects on Chikungunya transmission in Europe: geospatial analysis of vector’s climatic suitability and virus’ temperature requirements. Int J Health Geogr. 2013;12: 51.

30. Funk S, Kucharski AJ, Camacho A, Eggo RM, Yakob L, Murray LM, et al. Comparative analysis of dengue and Zika outbreaks reveals differences by setting and virus. PLoS Negl Trop Dis. 2016;10: e0005173.

31. Bastos L, Villela DA, Carvalho LM, Cruz OG, Gomes MF, Durovni B, et al. Zika in Rio de Janeiro: assessment of basic reproductive number and its comparison with dengue. BioRxiv. 2016; 055475.

32. Riou J, Poletto C, Boëlle P-Y. A comparative analysis of Chikungunya and Zika transmission. Epidemics. 2017;

33. Kearney M, Porter WP, Williams C, Ritchie S, Hoffmann AA. Integrating biophysical models and evolutionary theory to predict climatic impacts on species’ ranges: the dengue mosquito Aedes aegypti in Australia. Funct Ecol. 2009;23: 528–538.

34. Hopp MJ, Foley JA. Global-scale relationships between climate and the dengue fever vector, Aedes aegypti. Clim Change. 2001;48: 441–463.

35. Tesla B, Demakovsky LR, Mordecai EA, Ryan SJ, Bonds MH, Ngonghala CN, et al. Temperature drives Zika virus transmission: evidence from empirical and mathematical models. Proc R Soc Lond B Biol Sci. 2018;285: 20180795.

36. Hijmans RJ, Cameron SE, Parra JL, Jones PG, Jarvis A. Very high resolution interpolated climate surfaces for global land areas. Int J Climatol. 2005;25: 1965–1978.

37. Hijmans RJ, van Etten J. raster: Geographic analysis and modeling with raster data. [Internet]. 2012. Available: http://CRAN.R-project.org/package=raster

38. Center for International Earth Science Information Network (CIESIN), Columbia University. Gridded Population of the World, Version 4 (GPWv4). [Internet]. US NASA Socioeconomic Data and Applications Center (SEDAC); 2016. Available: http://sedac.ciesin.columbia.edu/data/set/gpw-v4-population-count-adjusted-to-2015-unwpp-country-totals

39. Moran AE, Oliver JT, Mirzaie M, Forouzanfar MH, Chilov M, Anderson L, et al. Assessing the global burden of ischemic heart disease: part 1: methods for a systematic review of the global epidemiology of ischemic heart disease in 1990 and 2010. Glob Heart. 2012;7: 315–329.

40. Ross N. fasterize: High performance raster conversion for modern spatial data. [Internet]. Available: https://github.com/ecohealthalliance/fasterize

41. Bhatt S, Gething PW, Brady OJ, Messina JP, Farlow AW, Moyes CL, et al. The global distribution and burden of dengue. Nature. 2013;496: 504–507.

42. Nsoesie EO, Kraemer M, Golding N, Pigott DM, Brady OJ, Moyes CL, et al. Global distribution and environmental suitability for chikungunya virus, 1952 to 2015. Euro Surveill Bull Eur Sur Mal Transm Eur Commun Dis Bull. 2016;21.

43. Samy AM, Thomas SM, Wahed AAE, Cohoon KP, Peterson AT. Mapping the global geographic potential of Zika virus spread. Mem Inst Oswaldo Cruz. 2016;111: 559–560.

44. Messina JP, Kraemer MU, Brady OJ, Pigott DM, Shearer FM, Weiss DJ, et al. Mapping global environmental suitability for Zika virus. Elife. 2016;5: e15272.

45. Bogoch II, Brady OJ, Kraemer M, German M, Creatore MI, Kulkarni MA, et al. Anticipating the international spread of Zika virus from Brazil. Lancet Lond Engl. 2016;387: 335–336.

46. Katzelnick LC, Gresh L, Halloran ME, Mercado JC, Kuan G, Gordon A, et al. Antibody-dependent enhancement of severe dengue disease in humans. Science. 2017;358: 929. doi:10.1126/science.aan6836

47. Salje H, Cummings DAT, Rodriguez-Barraquer I, Katzelnick LC, Lessler J, Klungthong C, et al. Reconstruction of antibody dynamics and infection histories to evaluate dengue risk. Nature. 2018;557: 719–723. doi:10.1038/s41586-018-0157-4

48. Flage R, Aven T. Emerging risk–Conceptual definition and a relation to black swan type of events. Reliab Eng Syst Saf. 2015;144: 61–67.

49. Musso D, Rodriguez-Morales AJ, Levi JE, Cao-Lormeau V-M, Gubler DJ. Unexpected outbreaks of arbovirus infections: lessons learned from the Pacific and tropical America. Lancet Infect Dis. 2018;

50. Ng V, Fazil A, Gachon P, Deuymes G, Radojević M, Mascarenhas M, et al. Assessment of the probability of autochthonous transmission of Chikungunya virus in Canada under recent and projected climate change. Environ Health Perspect. 2017;125.

51. Butterworth MK, Morin CW, Comrie AC. An analysis of the potential impact of climate change on dengue transmission in the southeastern United States. Environ Health Perspect. 2017;125: 579.

52. Acharya BK, Cao C, Xu M, Khanal L, Naeem S, Pandit S. Present and Future of Dengue Fever in Nepal: Mapping Climatic Suitability by Ecological Niche Model. Int J Environ Res Public Health. 2018;15: 187.

53. Equihua M, Ibáñez-Bernal S, Benítez G, Estrada-Contreras I, Sandoval-Ruiz CA, Mendoza-Palmero FS. Establishment of Aedes aegypti (L.) in mountainous regions in Mexico: increasing number of population at risk of mosquito-borne disease and future climate conditions. Acta Trop. 2017;166: 316–327.

54. Tjaden NB, Suk JE, Fischer D, Thomas SM, Beierkuhnlein C, Semenza JC. Modelling the effects of global climate change on Chikungunya transmission in the 21 st century. Sci Rep. 2017;7: 3813.

55. Tjaden NB, Caminade C, Beierkuhnlein C, Thomas SM. Mosquito-borne diseases: advances in modelling climate-change impacts. Trends Parasitol. 2017;

56. O’Neill BC, Done JM, Gettelman A, Lawrence P, Lehner F, Lamarque J-F, et al. The benefits of reduced anthropogenic climate change (BRACE): a synthesis. Clim Change. 2018;146: 287–301.

57. Lucey DR, Gostin LO. The emerging Zika pandemic: enhancing preparedness. Jama. 2016;315: 865–866.

58. Siraj AS, Perkins TA. Assessing the population at risk of Zika virus in Asia-is the emergency really over? BMJ Glob Health. 2017;2: e000309.

59. Shocket MS, Ryan SJ, Mordecai EA. Temperature explains broad patterns of Ross River virus transmission. Lipsitch M, editor. eLife. 2018;7: e37762. doi:10.7554/eLife.37762

60. Wesolowski A, Qureshi T, Boni MF, Sundsøy PR, Johansson MA, Rasheed SB, et al. Impact of human mobility on the emergence of dengue epidemics in Pakistan. Proc Natl Acad Sci. 2015; 201504964. doi:10.1073/pnas.1504964112

61. Liu-Helmersson J, Stenlund H, Wilder-Smith A, Rocklöv J. Vectorial Capacity of Aedes aegypti: Effects of Temperature and Implications for Global Dengue Epidemic Potential. PLoS ONE. 2014;9: e89783. doi:10.1371/journal.pone.0089783

62. Yang HM, Macoris MLG, Galvani KC, Andrighetti MTM, Wanderley DMV. Assessing the effects of temperature on the population of Aedes aegypti, the vector of dengue. Epidemiol Infect. 2009;137: 1188–1202. doi:10.1017/S0950268809002040

63. Hagel K, Milinski M, Marotzke J. The level of climate-change mitigation depends on how humans assess the risk arising from missing the 2 C target. Palgrave Commun. 2017;3: 17027.

64. Pongsiri MJ, Roman J, Ezenwa VO, Goldberg TL, Koren HS, Newbold SC, et al. Biodiversity loss affects global disease ecology. Bioscience. 2009;59: 945–954.

65. Zouache K, Fontaine A, Vega-Rua A, Mousson L, Thiberge J-M, Lourenco-De-Oliveira R, et al. Three-way interactions between mosquito population, viral strain and temperature underlying chikungunya virus transmission potential. Proc R Soc B Biol Sci. 2014;281: 20141078. doi:10.1098/rspb.2014.1078

66. Grau HR, Aide TM, Zimmerman JK, Thomlinson JR, Helmer E, Zou X. The ecological consequences of socioeconomic and land-use changes in postagriculture Puerto Rico. AIBS Bull. 2003;53: 1159–1168.

67. Li Y, Kamara F, Zhou G, Puthiyakunnon S, Li C, Liu Y, et al. Urbanization increases Aedes albopictus larval habitats and accelerates mosquito development and survivorship. PLoS Negl Trop Dis. 2014;8: e3301.

68. Ryan SJ, Ben-Horin T, Johnson LR. Malaria control and senescence: the importance of accounting for the pace and shape of aging in wild mosquitoes. Ecosphere. 2015;6: 170.

69. Tsetsarkin KA, Chen R, Weaver SC. Interspecies transmission and chikungunya virus emergence. Curr Opin Virol. 2016;16: 143–150.

70. Tsetsarkin KA, Chen R, Leal G, Forrester N, Higgs S, Huang J, et al. Chikungunya virus emergence is constrained in Asia by lineage-specific adaptive landscapes. Proc Natl Acad Sci. 2011; 201018344.

71. Moro ML, Gagliotti C, Silvi G, Angelini R, Sambri V, Rezza G, et al. Chikungunya virus in North-Eastern Italy: a seroprevalence survey. Am J Trop Med Hyg. 2010;82: 508–511.

72. Sissoko D, Ezzedine K, Moendandzé A, Giry C, Renault P, Malvy D. Field evaluation of clinical features during chikungunya outbreak in Mayotte, 2005–2006. Trop Med Int Health. 2010;15: 600–607.

73. Getz WM, Marshall CR, Carlson CJ, Giuggioli L, Ryan SJ, Romañach SS, et al. Making ecological models adequate. Ecol Lett. 2018;21: 153–166.

74. Petchey OL, Pontarp M, Massie TM, Kéfi S, Ozgul A, Weilenmann M, et al. The ecological forecast horizon, and examples of its uses and determinants. Ecol Lett. 2015;18: 597–611.

75. Beebe NW, Cooper RD, Mottram P, Sweeney AW. Australia’s dengue risk driven by human adaptation to climate change. PLoS Negl Trop Dis. 2009;3: e429.

76. Kraemer MU, Sinka ME, Duda KA, Mylne AQ, Shearer FM, Barker CM, et al. The global distribution of the arbovirus vectors Aedes aegypti and Ae. albopictus. eLife. 2015;4: e08347. doi:10.7554/eLife.08347

77. Brady OJ, Johansson MA, Guerra CA, Bhatt S, Golding N, Pigott DM, et al. Modelling adult Aedes aegypti and Aedes albopictus survival at different temperatures in laboratory and field settings. Parasit Vectors. 2013;6: 1–28. doi:10.1186/1756-3305-6-351

78. Benedict MQ, Levine RS, Hawley WA, Lounibos LP. Spread of The Tiger: Global Risk of Invasion by The Mosquito Aedes albopictus. Vector-Borne Zoonotic Dis. 2007;7: 76–85. doi:10.1089/vbz.2006.0562

79. Perkins TA, Siraj AS, Ruktanonchai CW, Kraemer MU, Tatem AJ. Model-based projections of Zika virus infections in childbearing women in the Americas. Nat Microbiol. 2016;1: 16126.

80. Carlson CJ, Dougherty E, Boots M, Getz W, Ryan S. Consensus and conflict among ecological forecasts of Zika virus outbreaks in the United States. Sci Rep. 2018;8: 4921.

81. Evans MV, Murdock CC, Drake JM. Anticipating Emerging Mosquito-borne Flaviviruses in the USA: What Comes after Zika? Trends Parasitol. 2018;

82. Olival K, Willoughby A. Prioritizing the “Dormant” Flaviviruses. EcoHealth. 2017;14: 1–2.

83. Gould EA, Higgs S. Impact of climate change and other factors on emerging arbovirus diseases. Trans R Soc Trop Med Hyg. 2009;103: 109–121.

